# The effect of prolonged elbow pain and rTMS on cortical inhibition: A TMS-EEG study

**DOI:** 10.1101/2024.11.26.625334

**Authors:** Nahian S Chowdhury, Wei-Ju Chang, Donovan Cheng, Naveen Manivasagan, David A Seminowicz, Siobhan M Schabrun

## Abstract

**Introduction:** Recent studies using combined transcranial magnetic stimulation (TMS) and electroencephalography (EEG) have shown that pain leads to an increase in the N45 peak of the TMS-evoked potential (TEP), which is mediated by GABAergic inhibition. Conversely, 10Hz repetitive TMS (10Hz-rTMS), which provides pain relief, reduces the N45 peak. However, these studies used brief pain stimuli (lasting minutes), limiting their clinical relevance. The present study determined the effect of pain and 10Hz-rTMS on the N45 peak in a prolonged pain model (lasting several days) induced by nerve growth factor (NGF) injection to the elbow muscle.

**Materials and Methods:** <u>Experiment 1</u>: TEPs were measured in 22 healthy participants on Day 0 (pre-NGF), Day 2 (peak pain), and Day 7 (pain resolution). <u>Experiment 2</u>: We examined the effect of 5 days of active (n=16) or sham (n=16) rTMS to the left primary motor cortex (M1) on the N45 peak during prolonged NGF-induced pain, with TEPs measured on Day 0 and Day 4 (post-rTMS).

**Results:** <u>Experiment 1:</u> While no overall change in the N45 peak was seen, a correlation emerged between higher pain severity on Day 2 and a larger increase in the N45 peak.

<u>Experiment 2</u>: Active rTMS reduced the N45 peak on Day 4 vs. Day 0, with no effect in the sham group.

**Conclusion:** Our findings suggest that (i) higher pain severity correlates with an increase in the N45 peak, and (ii) rTMS decreases cortical inhibition in a model of prolonged experimental pain. This study extends previous research by demonstrating a link between pain perception and cortical inhibition within a prolonged pain context.

## MAIN

The perception of pain arises from a complex interplay between emotional, cognitive and sensory mechanisms involved in the processing of afferent nociceptive inputs. Understanding these mechanisms is critical, as it can help us uncover pain biomarkers which can be applied to the diagnosis, prevention and treatment of chronic pain [1, 2]

Recent years have seen the use of combined transcranial magnetic stimulation and electroencephalography (TMS-EEG) to identify novel brain markers linked to pain and analgesia [3–6]. One of these markers is the TMS-evoked potential (TEP) N45 peak, which is usually observed ∼45ms after a TMS pulse is delivered to M1 [7]. The N45 peak is considered a measure of cortical inhibition, as its amplitude is respectively increased and decreased by GABA receptor agonists and antagonists [8–11]. In one study on healthy participants, across three experiments, thermal heat pain lasting ∼six minutes was shown to induce an increase in the N45 peak relative to a pain-free baseline, with larger increases in the N45 peak associated with greater pain sensitivity[3]. In another study on healthy participants, a single session of repetitive TMS (rTMS) to the posterior-superior insular cortex delivered during a capsaicin heat pain model lasting 90 minutes was shown to decrease heat pain sensitivity, with these analgesic effects accompanied by a decrease in the N45 peak. Moreover, compared to sham rTMS, the reductions in pain intensity following active rTMS were not just associated with, but partially mediated by, reductions in the N45 peak[4]. These findings suggest short lasting pain is associated with an increase in the N45 peak, while rTMS-induced analgesia is associated with a decrease in the N45 peak, supporting a link between pain perception and GABA-mediated cortical inhibition.

While these preliminary data show promise for the use of the N45 peak as a potential pain biomarker, past studies used short-lasting painful stimuli in the minutes range, which may have limited clinical relevance. It is unknown whether similar results are observed with longer lasting pain, such as pain induced by intramuscular injections of nerve growth factor (NGF) [12, 13]. NGF injections induce prolonged musculoskeletal pain [14] that mimics the duration, time course, functional limitation and hyperalgesia associated with chronic pain conditions [13, 15]. A single NGF injection to the elbow muscle induces pain mimicking lateral epicondylalgia over several days, with pain peaking at 1-2 days post-injection, and pain resolving ∼seven days post-injection[15]. Assessment of the effects of pain and rTMS on the TEPs during NGF-induced pain will therefore enhance our understanding of the cortical mechanisms implicated in pain and analgesia within a more clinically relevant prolonged pain experience.

The present study conducted two experiments which aimed to determine 1) the effect of prolonged lateral elbow pain induced by an NGF injection on cortical inhibition (assessed via the N45 peak), and 2) the effect of 5 consecutive days of rTMS on cortical inhibition within a model of prolonged lateral elbow pain. It was hypothesised that prolonged pain would increase the N45 peak relative to a pain-free baseline. Further, it was hypothesized that active rTMS would decrease the N45 peak relative to sham rTMS during pain.

## Methods

### Design

Both experiments were conducted at Neuroscience Research Australia Sydney. Experiment 1 was a longitudinal within-subjects design with participants followed for a period of 20 days. Experiment 2 was an assessor/participant blinded, randomised, sham-controlled, longitudinal, parallel design with participants followed for a period of 11 days. Experiment 2 was part of a larger study with three groups of participants: one group received individualized rTMS (where rTMS frequency was fixed to a person’s EEG peak alpha frequency), and the other two groups received either 10Hz or sham rTMS. Outcomes collected before and after rTMS were: pain, TMS-evoked potentials, corticomotor excitability and resting state EEG measures. Experiment 2 of this paper only reports the data for the effects of 10Hz and sham rTMS on TEPs, with the other outcomes to be reported elsewhere.

The primary outcome in both experiments was the TEP N45 peak, with pain being a secondary outcome. A detailed statistical analysis for the effects of rTMS on pain in Experiment 2 will be reported in a separate manuscript comparing individualized rTMS, 10Hz rTMS and sham rTMS. Nonetheless, in this paper, we report the effect size and Bayes factors comparing the max “pain at its worst” scores between active and sham rTMS groups to permit interpretation of the analgesic effects of rTMS. All procedures adhered to the Declaration of Helsinki, with written, informed consent obtained prior to study commencement. The study was approved by the local ethics committee at the University of New South Wales (HC230194 and HC230344).

### Sample Size Calculation

A sample size calculation was conducted (G*Power 3.1.9.7) for each experiment based on available means/SDs reported in previous studies assessing the effect of acute pain on the TEP N45 peak [3] and assessing the effect of 10Hz rTMS on the N45 peak [4] during pain. For Experiment 1, for the effect of pain on N45 peak (α = 0.05, β = 0.8, d = 0.75) a sample size of 16 was required to detect a within-group difference in the N45 peak between timepoints. We opted for a higher sample size of 22 participants to improve statistical power. For Experiment 2, for the effect of rTMS on the TEP N45 peak (α = 0.05, β = 0.8, effect size f = 0.31, correlation between repeated measures = 0.5) a sample size of at least 12 individuals was required to detect an interaction between time and group. We opted for a higher sample size of 16 participants in each group to improve statistical power.

### Participants

In Experiment 1, 22 healthy participants (11 females, 11 males, age: 22.0 ± 2.0) were recruited. In Experiment 2, 32 healthy participants (18 males, 14 females, age: 21.0 ± 2.0) were recruited. In both experiments, participants were included if they were between 18 and 65 years, and excluded if they presented with any acute pain, had a history or presence of chronic pain, neurological, musculoskeletal, psychiatric or other major medical condition, were pregnant and/or lactating, or were contraindicated for TMS (e.g., metal implants in the head) as assessed using the Transcranial Magnetic Stimulation Adult Safety Screen questionnaire [16].

### Protocol

Figure 1 shows the protocol for each experiment.

**Figure 1.**
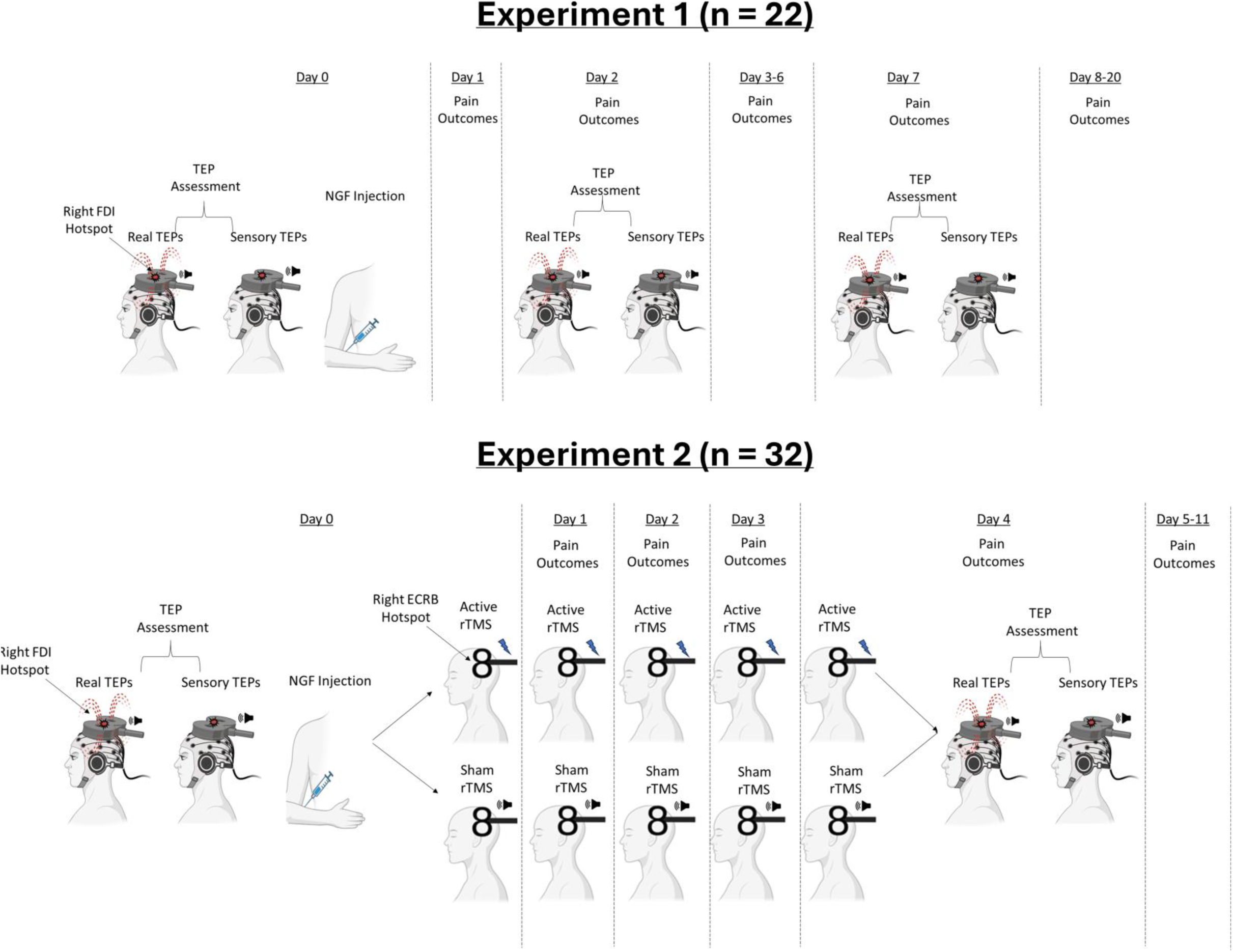
Diagram of the experimental protocols in each experiment.

#### Experiment 1

On Day 0, we first assessed “real” and “sensory” TEPs. Real TEP assessment involved TMS-EEG recording using a standard TMS coil. Sensory TEP assessment involved the use of a placebo coil which mimicked the auditory and somatosensory aspects of TMS, but without stimulating the brain. The rationale for using this coil is as follows: several studies have shown that a significant portion of TEPs do not reflect the direct cortical response to TMS, but rather auditory potentials elicited by the ‘clicking’ sound from the TMS coil, and somatosensory potentials elicited by the ‘flicking’ sensation on the skin of the scalp. This has led many authors to recommend the use of sensory control conditions to account for sensory contamination [17–20]. As such, sensory TEP assessment allowed us establish whether any changes in the N45 peak were related to the direct cortical response to TMS, rather than the sensory response to TMS. TEP measurement on Day 0 was followed by an NGF injection to the right ECRB muscle to induce prolonged elbow pain for several days. Questionnaires assessing pain, functional impairment and muscle soreness were sent to participant’s emails at 10am and 7pm each day from Days 1 to 20. TEPs were re-assessed on Day 2 (where pain was expected to peak) and Day 7 (where pain was expected to resolve) according to previous research[15].

#### Experiment 2

Prior to the study, each participant ID was randomly assigned to either an active or sham rTMS condition using a random number list generator (https://www.random.org/lists/). On Day 0, we assessed both real and sensory TEPs, followed by an NGF injection to the right ECRB. The allocations to either active or sham rTMS were then revealed to the experimenter (DC/NM) delivering rTMS via an automated email notification. The allocations were not revealed to the participant, or to the experimenters who collected TEP data or administered the injection of NGF (NC/WJC), and who performed data pre-processing and prepared the analysis plan (NC). Participants then received five daily sessions of 10Hz active or sham (audible clicks) rTMS each day from Day 0-4 by the same experimenter. The final rTMS session on Day 4 was followed immediately by assessment of real and sensory TEPs. Pain was assessed in the lab on Days 0 to 4, and via electronic diaries sent at 10am and 7pm each day from Days 5 to 11. Participants were debriefed (experimenter WJC) after completing the study (contacted by phone or email) and their group allocation was notified. Group allocations were revealed to experimenter NC when the data collection, pre-processing and analysis plan was complete.

### Data Collection Procedures

#### TEPs

Single, monophasic transcranial magnetic stimuli were delivered using a Duomag MP-EEG (Deymed, Czechia) and 70 mm figure-of-eight flat coil. EEG was recorded using a TMS-compatible amplifier (TruScan EEG 32, Deymed, Czechia) at a sampling rate of 3000 Hz. Signals were recorded from 32 passive electrodes, embedded in an elastic cap in line with the 10-5 system. Recordings were referenced online to ‘right mastoid’ and the ground electrode placed on the right cheekbone [4]. Electrolyte gel was used to reduce electrode impedances below ∼5 kΩ. In order to minimize the effect of the auditory response generated by the TMS coil click, a masking toolbox [66] was used with the participants wearing noise-cancelling headphones. In order to minimize the effect of the somatosensory response from the TMS coil, a thin layer of foam (5mm) was placed between the TMS coil and the scalp.

Surface disposable silver/silver chloride adhesive electrodes were applied over the right first dorsal interosseous (FDI) muscle. The coil was oriented at 45° to the midline, inducing a current in the posterior-anterior direction. To identify the left M1 target, the scalp site (‘hotspot’) that evoked the largest motor evoked potential (MEP) measured at the first dorsal interosseous (FDI) was determined and marked. The rest motor threshold (RMT) was determined using the ML-PEST (maximum likelihood strategy using parametric estimation by sequential testing) algorithm [21]. The test stimulus intensity was set at 90% RMT to minimize contamination of EEG signal from re-afferent muscle activation [22].

A real-time TEP visualization tool within the TruScan EEG software was used to confirm that artefacts (muscle, auditory) were minimal, and that, given the coil orientation and 90% RMT stimulus intensity, that early peaks (<100ms) at the stimulation site were evident (P30-N100) [6, 22, 23]. The visualization tool was used throughout each session to monitor coil positioning and TEP data quality across measurements within and between sessions. For each TEP measurement, 150 TMS pulses were delivered with a jitter of 1.75-2.25ms. TEPs were collected with both standard figure-of-eight coil to assess real TEPs (DuoMag 70 BF Coil) and a placebo coil to assess sensory TEPs (Duomag 70 BF Placebo coil). The placebo coil mimicked the sensations of real TMS (auditory and somatosensory) without stimulating the area under the coil. The order of the real and sensory TEP assessment was randomized for each session.

#### rTMS (Experiment 2 Only)

For the active rTMS group, 10Hz, biphasic TMS pulses were applied over the left primary motor cortex using a Magstim Super Rapid2 stimulator and a figure-of-eight coil. The location and intensity of rTMS was calibrated based on RMT of the right ECRB muscle (using the same hotspot and threshold procedures described previously). RMT was reassessed at each session to account for any changes in corticomotor excitability. During rTMS, the coil was held at 90° to the midline. 3000 stimuli 10Hz, 30 trains of 10 seconds, 20-second intertrain interval) were delivered at 90% of RMT. For sham rTMS, a sham coil that looked identical to the active rTMS but produced only audible clicks was used to deliver the stimulation protocol identical to the one used for active rTMS.

#### NGF Injection

A sterile solution of recombinant human NGF (dose of 5 μg [0.2 ml]) was administered as a bolus injection into the muscle belly of the ECRB using a 1-ml syringe with a disposable needle (27-G), retracted ∼2 mm.

#### Pain Outcomes

For Experiment 1, we used the PRTEE [24] Section 1 to assess pain while PRTEE Section 2 assessed functional impairment during specific and usual activities involving elbow muscles. Muscle soreness was assessed as described previously [12, 15].

Questionnaires were administered via automated links sent to participant’s email at 10am and 7pm each day. For Experiment 2, the PRTEE Section 1 was used assess pain in session on Days 0 to 4, and via electronic diaries at 10am and 7pm each day from Days 5 to 11.

### Data Processing

Pre-processing of the TEPs was completed using EEGLAB [25] and TESA [26] in MATLAB (R2021b, The Math works, USA), and based on previously described methods [26, 27]. First, the data was epoched 1000 ms before and after the TMS pulse, and baseline corrected between −1000 ms and −5 ms before the TMS pulse. The period between −5 ms and 15 ms after the TMS pulse was removed and interpolated by fitting a cubic function. Noisy epochs were identified via the EEGLAB auto-trial rejection function [28] and then visually confirmed. The fastICA algorithm with auto-component rejection was used to remove eyeblink and muscle artefacts [26]. The source-estimation noise-discarding (SOUND) algorithm was applied [27] to further supresses noise at each channel. The signal was then re-referenced (to average). A band-pass (1-100 Hz) and band-stop (48-52 Hz) Butterworth filter was then applied.

In line with previous studies investigating the effects of pain on TEPs [3], and rTMS on TEPs during pain [5], the average TEP was extracted from a-priori frontocentral region of interest (ROI) (’Fz’,’F3’,’F4’,’FC1’,’FC2’,’C3’,’Cz’, ‘C4’) investigated in previous studies [3, 4].

The N45 TEP peak from this ROI was identified for each participant using the TESA peak function [26], with a predetermined window of interest (40-60 ms) chosen to account for variation between participants in the latency of the peaks. Refer to the supplementary methods regarding analysis and results for the other peaks (N15, P30, P60, N100, P180).

### Statistical Analysis

Data are presented as mean ± standard deviations unless otherwise stated. Where violations of normality occurred according to Shapiro-Wilk tests, log-transformations of the data were conducted.

Bayesian inference was used to analyse the data, which considers the strength of the evidence for the alternative vs. null hypothesis, using JASP software (Version 0.12.2.0, JASP Team, 2020). Bayes factors were expressed as BF_10_ values, where BF_10_’s of 1–3, 3–10, 10– 30, 30-100 and >100 indicated ‘weak’, ‘moderate’, ‘strong’, ‘very strong’ and ‘extreme’ evidence for the alternative hypothesis, while BF_10_’s of 1/3–1, 1/10-1/3, 1/30-1/10 and 1/100-1/30 indicated ‘anecdotal’, ‘moderate’, ‘strong’, ‘very strong’ and ‘extreme’ evidence in favour of the null hypothesis. Default priors in JASP were used to provide a balance between informed and non-informed hypotheses.

#### Experiment 1

For the N45 peak, we conducted a 2(timepoint: Day 0, 2 and 7) x 2(stimulation: real vs. sensory) Bayesian Repeated measures ANOVA, followed by comparisons between Days 0 and 2, 0 and 7 and 2 and 7 using Bayes paired t-tests. An exploratory analysis on the association between N45 change at Day 2 for real TMS and mean pain, functional impairment and muscle soreness score on Day 2 (averaged across am and pm timepoints) was conducted.

#### Experiment 2

For the N45 peak, we conducted a 2(group: active vs. sham) 2x (stimulation: real vs. sensory) x 3(timepoint: Day 0, 2 and 7) Bayesian mixed-model ANOVA. We then conducted two 2-way Bayesian ANOVAs for real and sensory TEPs separately, followed by comparisons (Bayes t-tests) between Days 0 and Day 4 for the active and sham rTMS groups separately.

## Results

### Experiment 1 Results

All participants attended all three sessions with no missing data. The NGF injection led to no side effects. The mean (±SD) RMT on Day 0, Day 2 and Day 7 was 57.3 ± 7.7, 57.7 ± 8.2 and 56.2 ± 7.8 respectively.

Grand-average TEPs and scalp topographies for Day 0, 2 and 5 for real and sensory TEPs are shown in Figures 2 and 3 respectively. Figure 4 shows the grand-averages for the frontocentral ROI at Day 0, 2 and 5 for real and sensory TEPs. A two-way Bayesian repeated measures ANOVA revealed very strong evidence that the N45 peak was higher for real vs. sensory TEPs (BF_10_=91.88), moderate evidence for no difference in the N45 peak between days (BF_10_=0.33) and moderate evidence for no interaction between time and stimulation (BF_10_=0.25). When analysing the real TEPs alone, Bayesian t-tests revealed moderate evidence for no difference in the N45 peak between Days 0 and Day 2 (BF_10_=0.22), anecdotal evidence for no difference between Days 0 and 7 (BF_10_=0.83), and anecdotal evidence for a decrease in the N45 peak from Day 2 to Day 7 (BF_10_=2.00).

**Figure 2.**
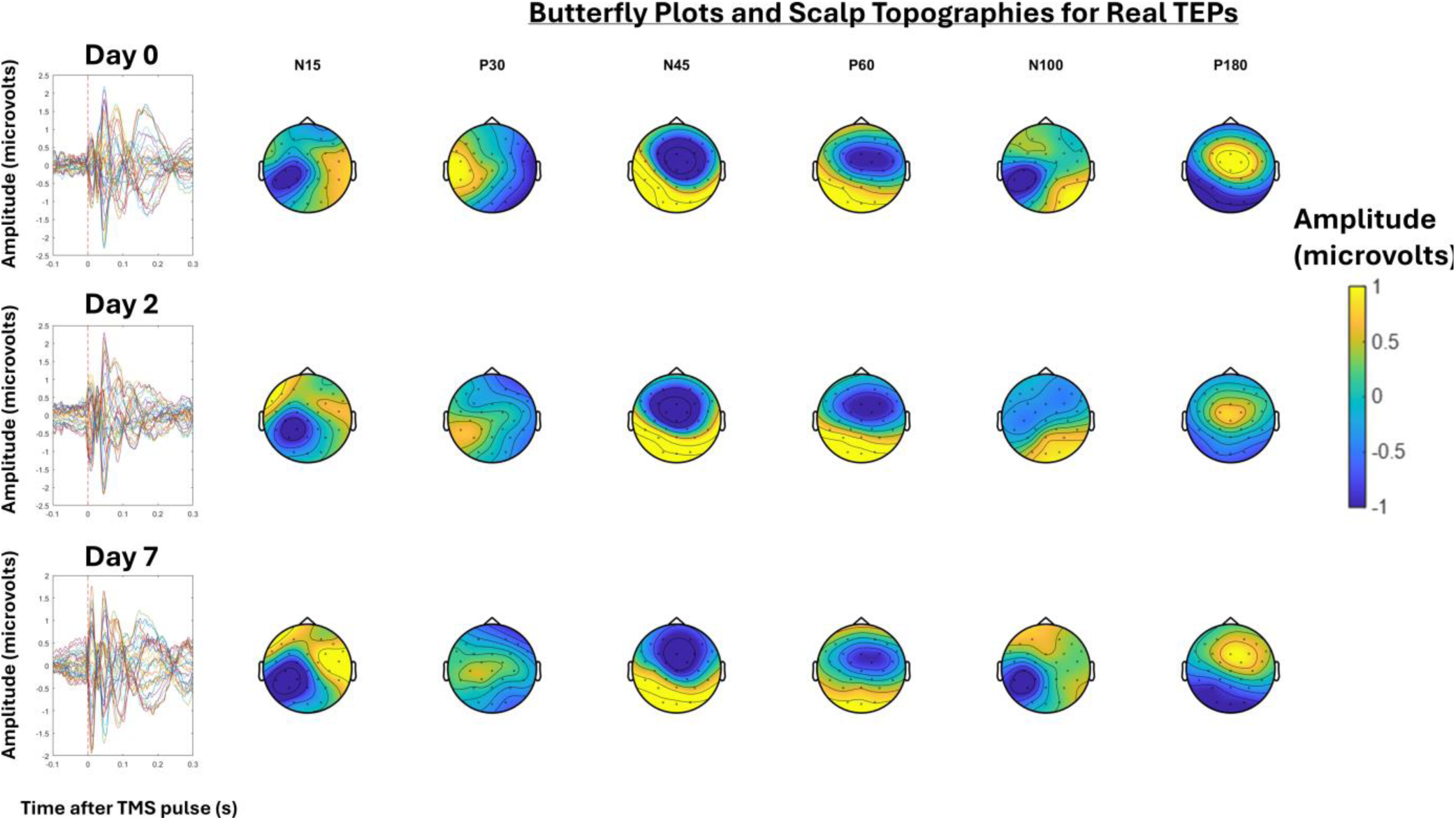
The Grand-average TEPs (n = 22) for all channels for the “real TEP” assessment at Day 0, 2 and 7. Scalp topographies and estimated source activity at timepoints where TEP peaks are commonly observed, including the N15, P30, N45, P60, N100, and P180.

**Figure 3.**
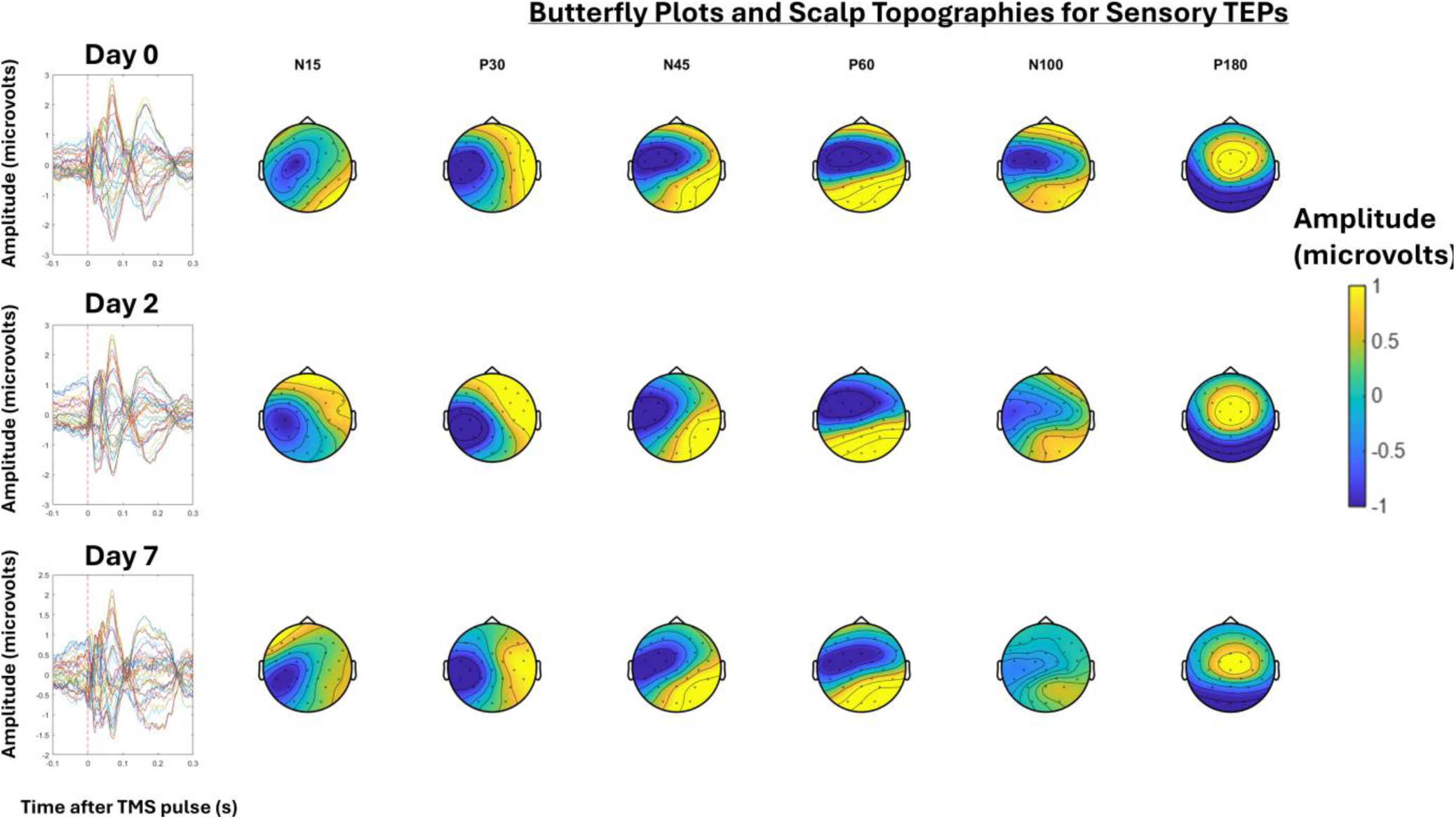
The Grand-average TEPs (n = 22) for all channels for the “sensory TEP” assessment at Day 0, 2 and 7. Scalp topographies and estimated source activity at timepoints where TEP peaks are commonly observed, including the N15, P30, N45, P60, N100, and P180.

**Figure 4.**
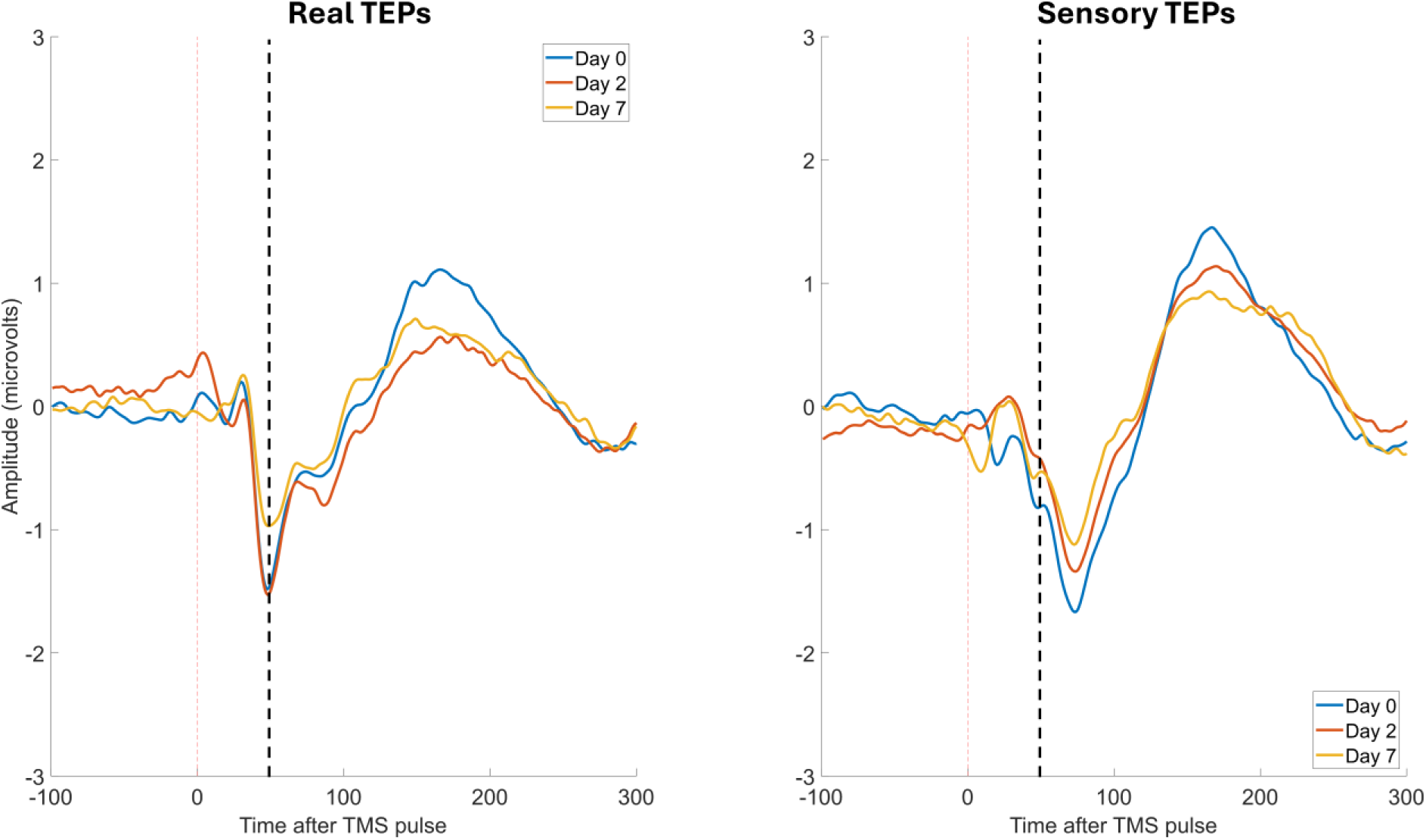
Prolonged pain did not alter the TEP N45 peak. The Grand-average TEPs (n=22) for the frontocentral ROI for “real TEPs” assessment on the left and “sensory TEPs” assessment on the right, across Day 0, 2 and 7. The black dotted line shows the approximate timing of the N45 peak.

Figure 5 shows the mean PRTEE and muscle soreness scores across Days 0-20. The mean maximum scores for PRTEE, pain at its worst, functional impairment and muscle soreness were respectively 7/50 ± 5.9, 2.2/10 ± 1.6, 9.2/100 ±10.4 and 2.6/7 ± 1.5. This indicates that, at its worst, participants experienced mild-moderate pain, functional impairment and muscle soreness. Figure 5 also shows the individual level relationships between pain intensity, functional impairment and muscle soreness on Day 2, and the increase in the N45 for both real and sensory TEPs measured on Day 2. For real TEPs, there was moderate evidence that a larger increase in the N45 peak was correlated with higher pain (r=-.56, BF_10_=7.9), functional impairment, (r=-0.53, BF_10_=5.5) and muscle soreness (r=-0.51, BF_10_=4.10). For sensory TEPs, there was anecdotal evidence for no relationship between changes in the N45 peak and pain (r=0.33, BF_10_=0.73) functional impairment (r=.18, BF_10_=0.36) and muscle soreness (r=0.32, BF_10_=0.71).

**Figure 5.**
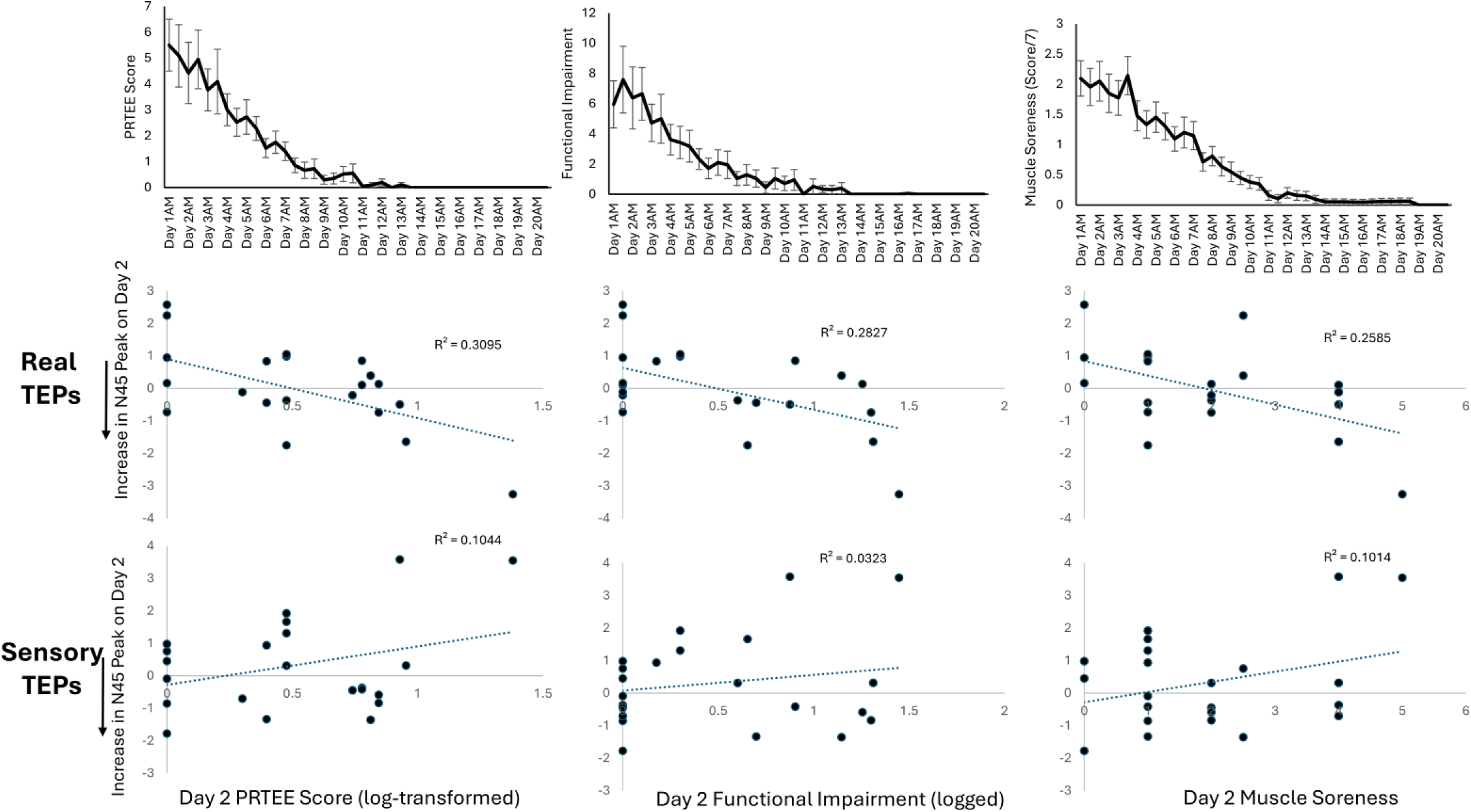
A larger increase in the TEP N45 peak was associated with higher pain, functional impairment and muscle soreness on Day 2. Panel A shows the mean (±SEM) PRTEE, functional impairment and muscle soreness scores across the 20 day period. Panel B shows the relationship between the change in the TEP N45 peak at Day 2 and PRTEE score, functional impairment score and muscle soreness score on Day 2 (averaged across am and pm timepoints).

### Experiment 2 Results

All 32 participants attended the 5 lab sessions with no missing data. All participants tolerated the rTMS and NGF injection without side effects. The mean RMTs for the active rTMS group was 59.2 ± 6.0 and 59.3 ± 7.1 on Days 0 and 4 respectively. The mean RMTs for the sham rTMS group was 62.2 ± 7.9 and 63.9 ± 8.9 on Days 0 and 4 respectively.

Figure 6 shows the grand-average TEPs and scalp topographies at Day 0 and Day 4 for the active rTMS group, for both real and sensory TEPs. Figure 7 shows the grand-average TEPs and scalp topographies at Day 0 and Day 4 for the sham rTMS group, for both real and sensory TEPs. Figure 8A shows the PRTEE scores for the active and sham rTMS group across Days 0-11. For the active rTMS group, the mean maximum PRTEE and “pain at its worst” score was 5.5/50 ± 3.3 and 2.0/10 ± 1.3. For the sham rTMS group, the mean maximum PRTEE and “pain at its worst” score was 7.3/50 ± 5.5 and 2.5/10 ± 2.0 respectively. The effect sizes (Cohen’s d) for the effect of rTMS on worst PRTEE score and pain at its worst were respectively 0.39 and 0.3, indicating a small-medium effect of active rTMS on pain relative to sham rTMS. However, there was anecdotal Bayesian evidence for no difference in maximum PRTEE (BF_10_=0.5) and “pain at its worst” (BF_10_=0.4) scores between active and sham rTMS groups.

**Figure 6.**
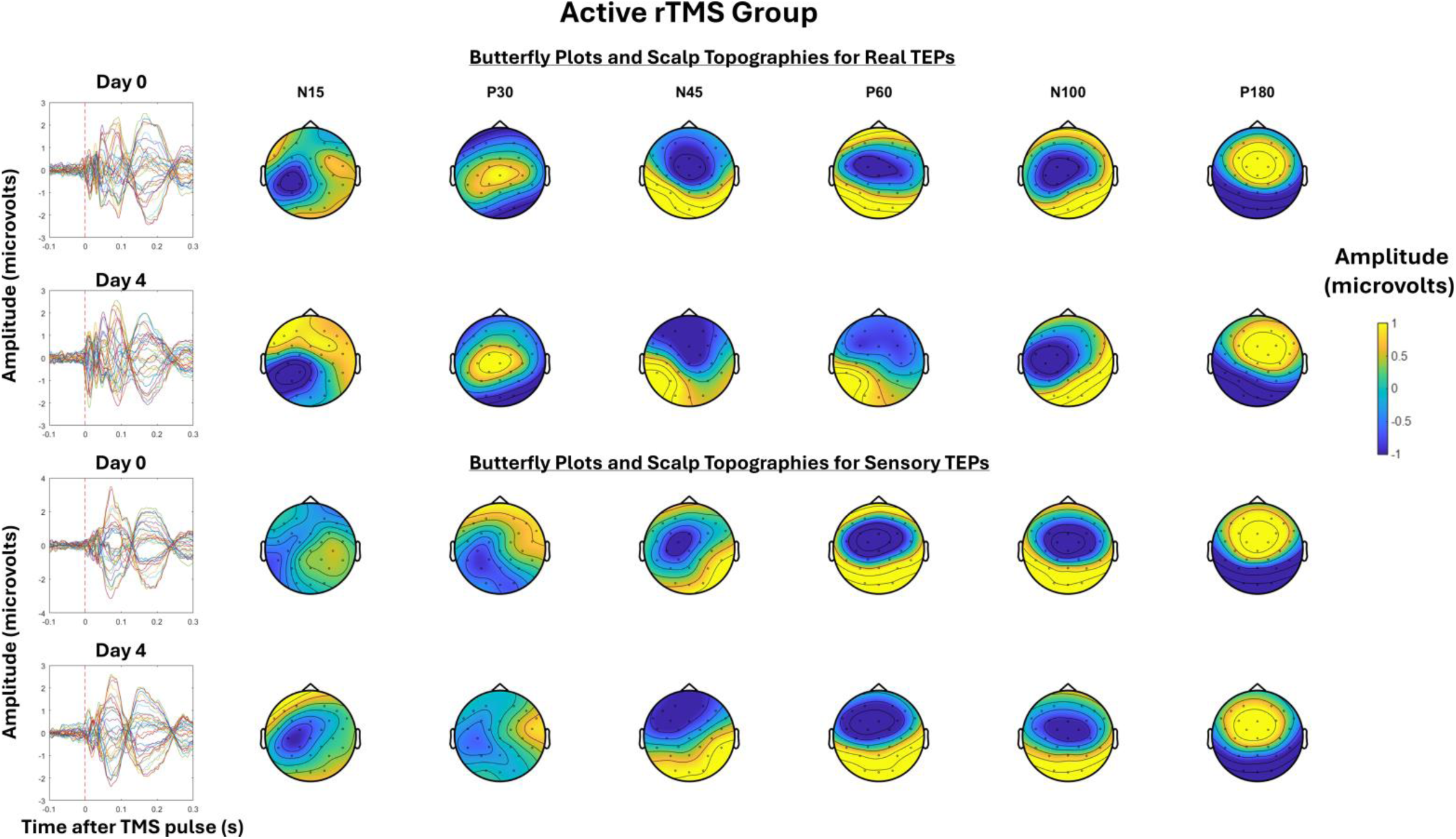
The left shows the Grand-average TEPs (n = 16) for all channels for the TEP assessment at Day 0 and 4 for the active rTMS group. The right shows the scalp topographies and estimated source activity at timepoints where TEP peaks are commonly observed, including the N15, P30, N45, P60, N100, and P180. The top half shows the real TEP waveforms and scalp topographies, while the bottom half shows the sensory TEP waveforms and scalp topographies.

**Figure 7.**
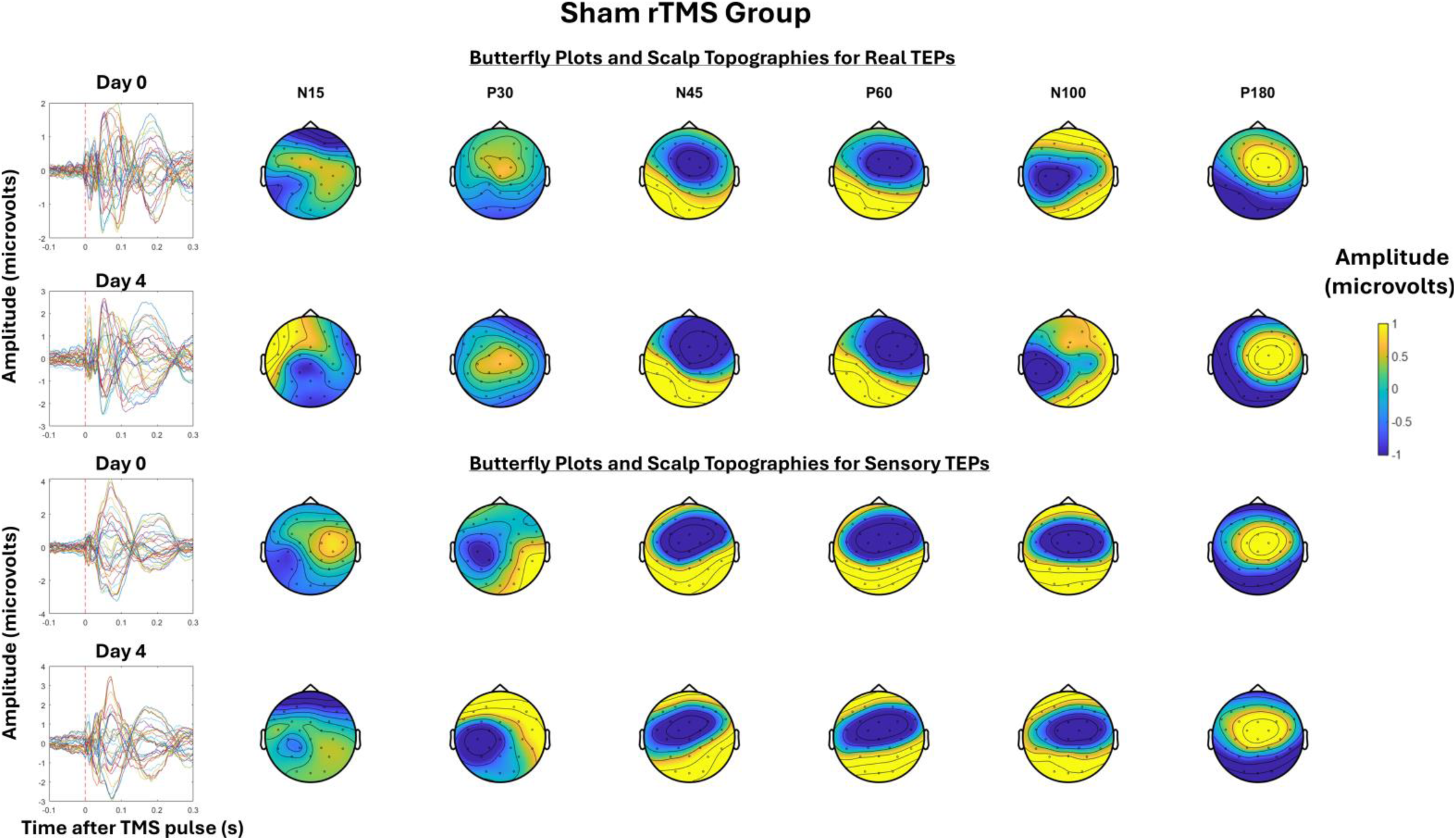
The left shows the Grand-average TEPs (n = 16) for all channels for the TEP assessment at Day 0 and 4 for the sham rTMS group. The right shows the scalp topographies and estimated source activity at timepoints where TEP peaks are commonly observed, including the N15, P30, N45, P60, N100, and P180. The top half shows the real TEP waveforms and scalp topographies, while the bottom half shows the sensory TEP waveforms and scalp topographies.

**Figure 8.**
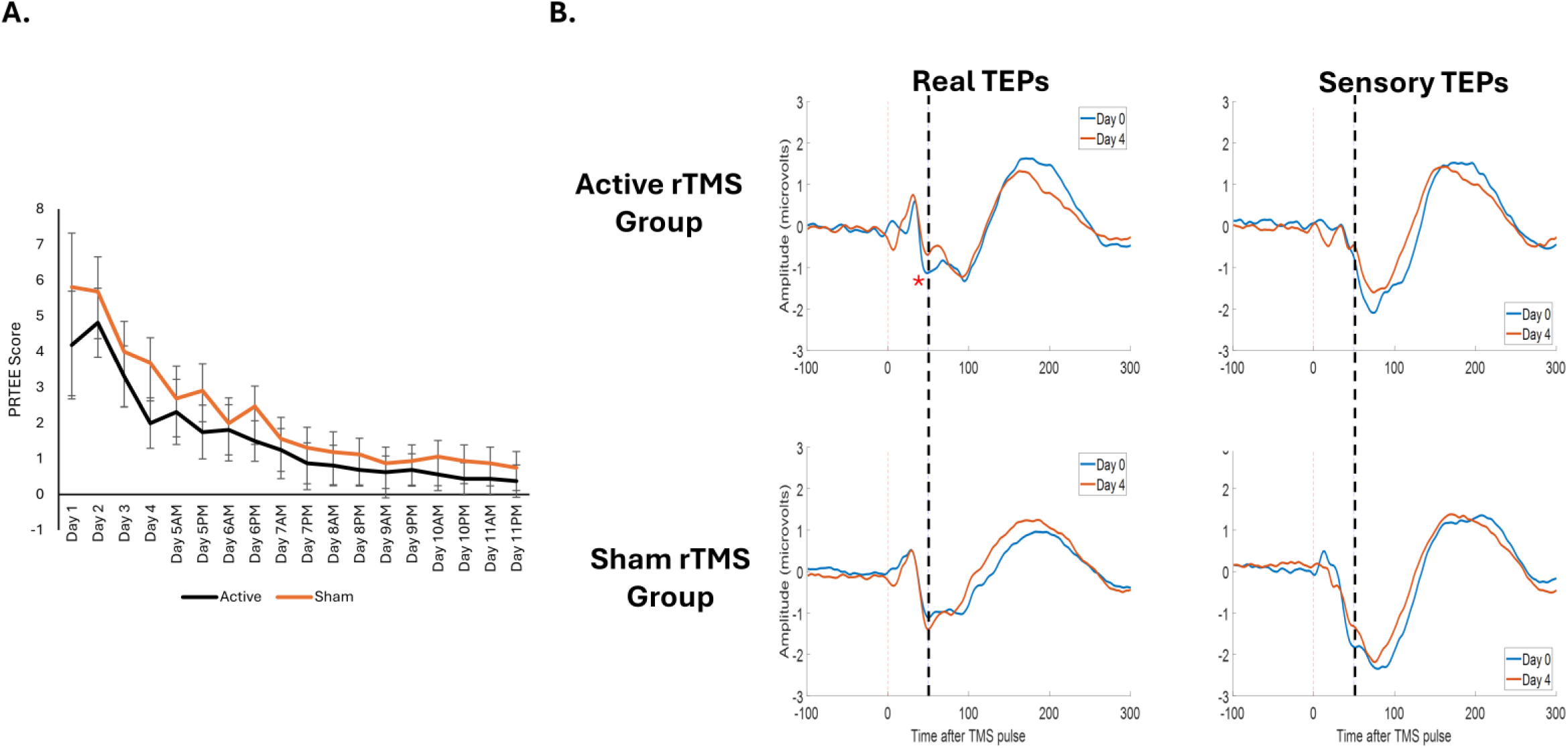
5 days of Active rTMS led to a decrease in the TEP N45 peak. The left shows the mean (±SEM) PRTEE scores for the active and sham rTMS groups across the 11 day period. The right shows the grand-average TEPs for the frontocentral ROI for the active rTMS group (left, n = 16) and sham rTMS group (right, n = 16) on the right, with waveforms for real TEPs on the upper panel and waveforms for sensory TEPs on the lower panel. The blue dotted line shows the approximate timing of the N45 peak.

Figure 8B shows the grand-average TEPs for the frontocentral ROI at Days 0 and 4 for real and sensory TEPs across active and sham rTMS groups. A three-way Bayesian mixed-model ANOVA revealed moderate evidence for a three-way interaction between stimulation, group and time, such that, relative to sham, the decrease in the N45 peak following 5 days of active rTMS was stronger for real TEPs compared to sensory TEPs (BF_10_=6.5). When analysing real TEPs alone, there was moderate evidence for a group x time intervention, such that the decrease in the N45 peak was larger following 5 days of active rTMS compared to sham rTMS (BF_10_=7.8). Bayesian paired t-tests showed strong evidence that the N45 peak decreased from Day 0 to Day 4 in the active rTMS group (BF_10_=13.45), and anecdotal evidence for no change in the N45 peak in the sham Rtms group (BF_10_=0.4). For sensory TEPs, there was anecdotal evidence for no group x time intervention, such that relative to sham rTMS, active rTMS did not modulate the N45 peak of sensory TEPs (BF_10_=0.45).

## Discussion

The present study aimed to determine the effects of pain and rTMS on cortical inhibition (indexed via the N45 peak) within a model of prolonged lateral elbow pain induced by an NGF injection. Contrary to our hypotheses, we did not find a group level effect of prolonged pain on cortical inhibition in Experiment 1. However, exploratory analysis showed that an increase in cortical inhibition was associated with higher pain intensity. Experiment 2 showed that, consistent with our hypothesis, 5 days of 10Hz rTMS over M1 led to a decrease in cortical inhibition during prolonged lateral elbow pain. Here we discuss these results in detail.

### Higher pain is associated with an increase in the TEP N45

Experiment 1 examined the effect of prolonged pain on the N45 peak. Against predictions, prolonged pain did not induce a group level effect on the N45 peak. This is inconsistent with our previous work that showed that acute heat pain lasting several minutes led to an increase in the N45 peak [3]. One explanation for the lack of an effect is that pain and muscle soreness induced by the single injection to the ECRB was on average, mild to moderate (worst pain and muscle soreness was 2.3/10 and 2.7/7 respectively). Nonetheless, we found that the increase in the N45 peak was correlated with higher pain, which is consistent with a previous study showing heat pain severity correlated with the increase in the N45 peak [3]. As such, the NGF model is likely more suited for understanding individual differences in pain sensitivity, rather than the group effect of pain. This is consistent with literature exploring the effect of NGF pain on corticomotor excitability. In a systematic review [29], it was shown that prolonged pain (in the days-week range) did not decease corticomotor excitability on a group level, but that larger decreases in corticomotor excitability were correlated with higher pain severity. It is plausible that when pain lasts only seconds to minutes, this is an immediate threat to the affected area, resulting in more uniform changes in cortical excitability. However, when pain lasts for days or weeks, how each individual responds to pain and how brain regions involved in pain processing function vary. For example, in some individuals pain processing is more inhibited to prioritise task completion, while for others, pain processing dominates over other functions [30–32]. These differing pain adaptation strategies are associated with substantial variations in the brain’s response to pain over time [33, 34]. Overall, our results provide further support for the role of N45 peak within a more clinically relevant prolonged pain experience, necessitating further investigation in clinical pain populations.

### While rTMS did not reduce pain, it led to a reduction in the N45 peak

We found that 5 days of active rTMS decreased the N45 peak, with no changes in the N45 peak following sham rTMS. These findings are consistent with our previous study showing that a single session of 10Hz rTMS to the posterior superior insular cortex led to a reduction in the N45 peak during pain [4], suggesting that the effects of rTMS on the N45 peak generalized across different stimulation sites. These findings are also consistent with other research showing decreases in the N45 peak following multiple sessions of 10Hz rTMS to the dorsolateral prefrontal cortex in patients with depressive symptoms [35]. As we did not assess the N45 peak in response to pain alone (since the rTMS was given immediately after the NGF injection), it was not possible to determine whether any reductions in pain from pre to post-rTMS were correlated with, or mediated by, the reductions in the N45 peak induced by rTMS. Nonetheless, our findings suggest that rTMS can reduce the N45 peak in people experiencing prolonged experimental pain. We encourage future larger scale studies to determine whether interventions that can specifically target the N45 peak can also induce larger reductions in pain.

### Strengths, Limitations and Future Suggestions

A strength of the study is the use of a sensory control condition for the TEP measurements. Previous studies have shown that even when masking methods are used to minimize auditory or somatosensory effects of TMS, sensory contamination of TEPs is still present [20], as shown by strong temporal and spatial correlations in the signals of real TMS and sensory control conditions that mimic the auditory/somatosensory aspects of real TMS [17, 19]. This has led to recommendations to use sensory control conditions [17, 19].

However many authors opt not to use a sensory control condition due to the additional testing times and/or lack of resources to develop sensory control conditions (e.g. absence of a sham coil or sufficient apparatus to replicate auditory and somatosensory aspects of real TMS). Here, in both experiments, we included a sensory control condition using a coil (Duomag 70BF placebo coil) that could not only generate a click noise, but which could also induced a somatosensory response (flick sensation), which most commercially available sham coils are unable to achieve. We showed that the relationship between pain and the N45 peak occurred only for real and not sensory TEPs (Experiment 1), and that active rTMS only modulated real and not sensory TEPs (Experiment 2), thus strengthening our conclusions.

Despite the abovementioned strength, the main limitation across both experiments was the use of a single NGF injection, inducing a mild pain that may not have been sufficient to induce an effect on the N45 peak in Experiment 1, and may also reduced the magnitude of any rTMS-induced analgesic effects in Experiment 2. Future research should consider using multiple NGF injections to induce pain with higher severity and longer duration. Another limitation is the use of a 32-channel EEG montage. Although 32 channels can be used to reliably obtain TEPs [5, 36, 37], a higher electrode array could provide a higher spatial resolution of the TEP response to pain and rTMS. Nonetheless, the electrodes analysed in both experiments were consistent with ROI from previous studies [3, 4], with these electrodes demonstrating similar changes in response to pain and rTMS. Lastly, while the inclusion of a sensory control condition was a strength, the N100 peak of the sensory TEPs was quite large relative to real TEPs, possibly due to the louder click sound generated from the sham coil, even at the same stimulation intensity as the active coil [38]. Future studies are encouraged to adjust for the sound intensity of the sham coil so that it matches the intensity in the active coil.

### Conclusion

Our study showed that while prolonged experimental pain did not change the N45 peak, higher pain severity was associated with a larger increase in the N45 peak. Moreover, 5 daily sessions of 10Hz rTMS over M1 led to a reduction in the N45 peak during prolonged experimental pain. This study provides further evidence for a link between cortical inhibition and pain perception, showing that higher pain severity is associated with an increase in the N45 peak, and rTMS decreases cortical inhibition during prolonged experimental pain. Our findings extend previous research beyond transient painful stimuli to a model of prolonged pain and provide further evidence for a link between pain perception and cortical inhibition.

## Author Contributions

**Nahian S Chowdhury:** Conceptualization, Data curation, Formal Analysis, Funding acquisition, Investigation, Methodology, Project Administration, Resources, Software, Supervision, Validation, Visualization, Writing – Original Draft, Writing – Reviewing and Editing. **Wei-Ju Chang**: Conceptualization, Data curation, Formal Analysis, Investigation, Methodology, Project Administration, Writing – Original Draft, Writing – Reviewing and Editing. **Donovan Cheng:** Investigation, Methodology. **Naveen Manivasagan:** Investigation, Methodology. **David Seminowicz:** Funding acquisition, Project Administration, Resources, Software, Supervision and Writing – Reviewing and Editing. **Siobhan M Schabrun:** Conceptualization, Funding acquisition, Project Administration, Resources, Software, Supervision and Writing – Reviewing and Editing.

