## Supplementary Material for "The effect of prolonged elbow pain and rTMS on cortical inhibition: A TMS-EEG study"

^1^Center for Pain IMPACT, Neuroscience Research Australia, Sydney, New South Wales, Australia, ^2^University of New South Wales, Sydney, New South Wales, Australia, ^4^Department of Medical Biophysics, Schulich School of Medicine & Dentistry, University of Western Ontario, London, Canada, ^5^The Gray Centre for Mobility and Activity, Parkwood Institute, St. Joseph’s Healthcare, London, Canada, ^6^School of Physical Therapy, University of Western Ontario, London, Canada

**Supplementary Methods**

**Additional Peak Analysis**

Other peaks of the TEPs from the frontocentral ROI (N15, P30, P60, N100, P180) were identified for each participant using the TESA peak function, with predetermined windows of interest (N15: 12-20 ms, P30: 25-40 ms, P60: 55-70 ms, N100: 70-110 ms, P180: 150-200 ms). The same statistical analysis as with the N45 peak was conducted on these other peaks.

**Supplementary Results**

**Experiment 1 Other Peaks**

Supplementary Table 1. Bayes Factors assessing the difference in peak amplitude

| Peak | Day 0 vs. Day 2 (Real TEPs) | Day 0 vs. Day 7  (Real TEPs) | Day 2 vs. Day 7  (Real TEPs) |
| --- | --- | --- | --- |
| N15 | 0.23 | 0.27 | 0.23 |
| P30 | 0.26 | 0.22 | 0.35 |
| P60 | 0.23 | 0.27 | 0.27 |
| N100 | 0.28 | 0.29 | 0.52 |
| P180 | 1.8 | 1.3 | 0.24 |

**Experiment 2**

Supplementary Table

| Peak | Time x Group Interaction for Real TEPs |
| --- | --- |
| N15 | 0.798 |
| P30 | 0.266 |
| P60 | 0.52 |
| N100 | 0.447 |
| P180 | 4.3  Active rTMS (Day 0 vs. Day 4): 2.54  Sham rTMS (Day 0 vs. Day 4): 0.46 |
